# Regulatory network architecture constrains inflammatory responses in tissue-resident alveolar macrophages

**DOI:** 10.64898/2026.02.16.706134

**Authors:** Sonia M. Kruszelnicki, Sreeparna Chakraborty, Xinge Wang, Jalees Rehman, Harinder Singh, Rachel A. Gottschalk

## Abstract

Macrophages across different tissues exhibit remarkable functional diversity while maintaining core innate immune-cell programming. These complex programs are governed by gene regulatory networks, in which precise transcription factor activity tunes the expression of functionally relevant gene modules. Although the contributions of individual transcription factors have been well characterized, the higher-order regulatory interactions that coordinate tissue-resident macrophage identity and inflammatory response regulation remain poorly understood. Here, we integrate single-cell RNA-seq data with ATAC-seq profiling and deep-learning-based chromatin accessibility modeling to infer gene regulatory network architectures in tissue-resident versus recruited monocyte-derived alveolar macrophages under inflammatory stress. Our results suggest that inflammatory responses are more restrained in tissue-resident alveolar macrophages compared with recruited macrophages due to a stabilizing regulatory network architecture involving PU.1 and CEBP/β. This work advances our understanding of functional plasticity in tissue-resident macrophages and their role in host defense.

## Introduction

Tissue-resident alveolar macrophages (AMs) have specialized homeostatic functions, including catabolism of lipid-rich pulmonary surfactant to support gas exchange and clearance of inhaled debris. These functions are programmed by the expression and activities of transcription factors (TFs) and stimuli specific to the alveolar compartment of the lung, such as GM-CSF(1) and TGF-β(2) (3, 4). Over the course of inflammation, AMs shift from a homeostatic, tissue-maintenance state to a pro-inflammatory state, followed by states associated with resolution and tissue repair, before ultimately returning to homeostasis. Dysregulated macrophage states have been associated with chronic diseases, aging, microbial dysbiosis, and environmental pollution(5–9), highlighting the importance of tight regulatory control over macrophage transcriptional state to maintain healthy tissue.

Context-dependent macrophage states are governed by TFs that regulate functionally relevant gene expression programs and comprise key nodes within gene regulatory networks (GRNs). Such networks provide comprehensive frameworks for analyzing and modeling TF-target gene interactions(10–12). While previous studies have constructed inflammatory response GRNs using *in vitro* macrophage systems(13, 14), such GRNs did not address how tissue-specific TF activity intersects with inflammatory response regulation in vivo. Here, we take an integrative approach to examine regulatory network structures that coordinate AM functional programs across homeostasis and inflammation. Considering the known differences in transcriptional and chromatin landscapes between tissue resident and recruited AMs (15), we assembled GRNs specific to each group to determine how TFs that maintain tissue-resident AM identity shape their inflammatory response regulation. We apply topic modeling to define distinct functional programs and then link these programs in turn to their transcriptional regulators via the assembled GRNs. Our findings suggest that despite experiencing the same environmental signals, inflammatory responses are more restrained in tissue-resident AMs than in recruited monocyte-derived AMs, and that this restraint is maintained by a stabilizing GRN structure that preserves tissue integrity.

## Results and Discussion

To delineate dynamically regulated gene expression programs and assemble the underlying GRNs controlling AM responses to acute lung inflammation, we analyzed a recently published murine scRNA-seq dataset (16). Immediately after euthanasia, anti-mouse CD45 antibody was instilled intratracheally to label cells in the airspace and macrophages (CD88+CD64+ and i.t. anti-CD45+) were isolated and profiled at steady state and at 3, 6, and 15 days following intratracheal administration of LPS(16) (Figure 1A). We applied topic modeling to identify major gene expression programs across the LPS stimulation time course. Using grade-of-membership analysis(17), we performed KEGG pathway enrichment on the top 1,000 genes for each topic to functionally annotate topics as states of homeostasis, inflammation, or resolution(18)(Figure 1B,C). The inflammation topic was highly enriched for pathways associated with disease states and pro-inflammatory signaling, while the homeostasis topic was enriched for terms related to cell cycle regulation and lipid processing, including the PPAR signaling pathway, consistent with the established role of PPAR-γ in coordinating AM surfactant homeostasis (19, 20). We next annotated cells as tissue-resident or recruited AMs based on k-means clustering and expression of tissue-resident and recruited, monocyte-derived AM marker genes (Figure 1D,E). As expected, recruited AMs were largely represented during inflammation and resolution, at days 3 and 6 post-LPS, respectively. In contrast, tissue-resident AMs were more evenly distributed across timepoints, with cells manifesting low levels of the inflammation topic, and instead remaining associated with the homeostasis gene topic despite LPS challenge. Together, these analyses, based on linking topic-based gene expression programs with AM functional states and identities, highlighted a major difference in the inflammatory response of tissue resident versus recruited AMs.

**Figure 1.**
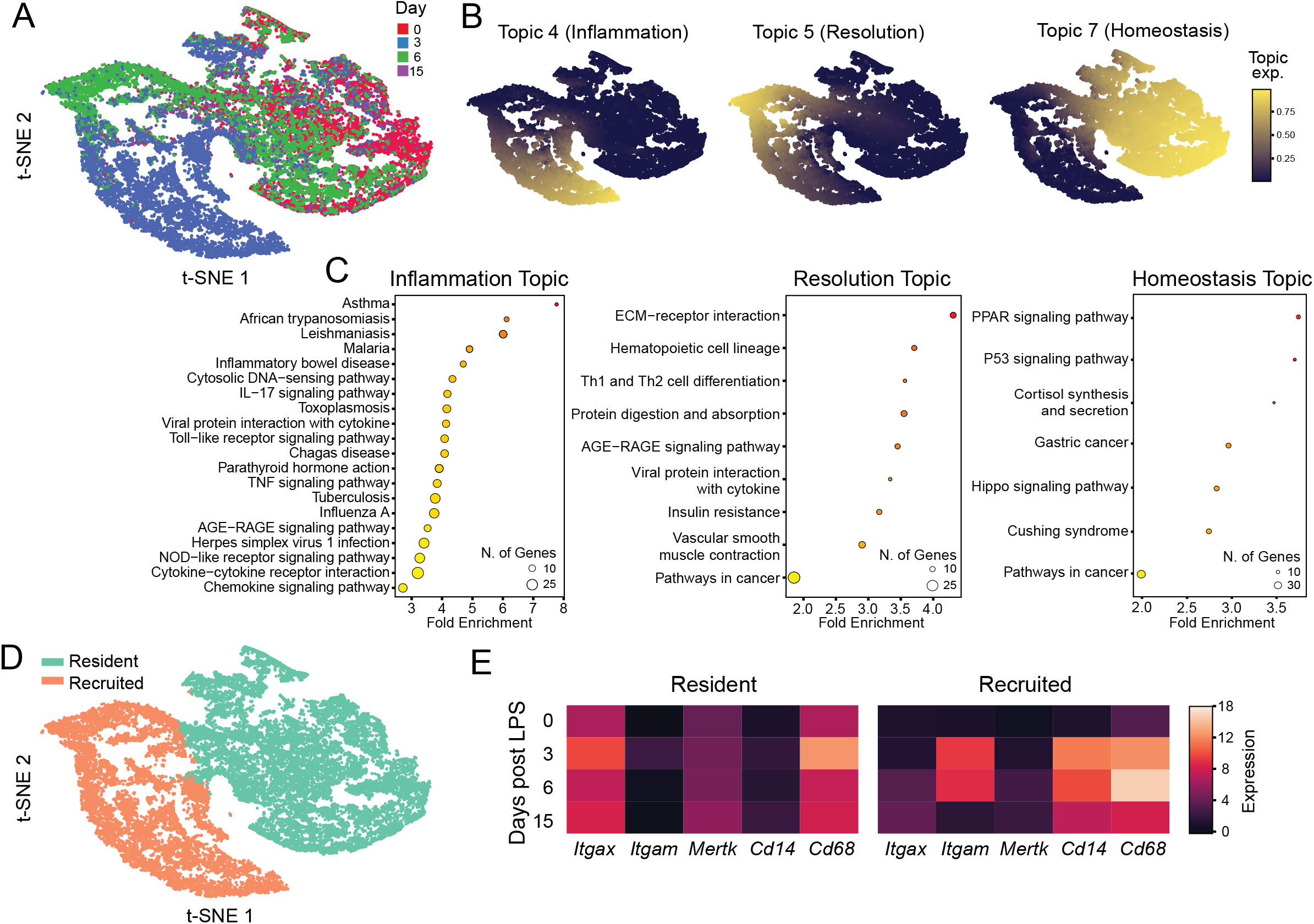
Defining alveolar macrophage gene topics associated with homeostasis and inflammation. Processed scRNA-seq data from King et al. of sorted AMs (CD88+CD64+ and i.t. anti-CD45+) were isolated at 0, 3, 6, and 15 days following intratracheal administration of LPS. t-SNE plots colored by (a) timepoint after LPS-induced inflammation and (b) by topic-expression as determined by the loadings matrix of the multinomial topic model. c) Enriched KEGG pathways for top 1000 differentially expressed genes for inflammation, resolution, and homeostasis topics 4, 5, and 7, respectively. Circle size corresponds to the number of terms in the pathway and color corresponds to significance of enrichment. d) t-SNE plot showing the clusters annotated as recruited and resident AMs e) heatmaps of pseudobulked expression for resident and recruited AMs of characteristic genes at each LPS timepoint.

We next sought to understand the gene regulatory architectures governing these distinct inflammatory AM responses. We used CellOracle(12) to build GRNs specific to tissue-resident or recruited AMs. Across the different timepoints following LPS administration, TFs in the tissue-resident population exhibited higher average eigenvector centrality within their GRNs compared to TFs in recruited AM GRNs (Figure 2A). Eigenvector centrality reflects the extent to which a TF is connected to other highly connected nodes, which in this context are predominantly other TFs. Reasoning that increased TF-TF connectivity would confer greater redundancy and stability of regulatory activity, we systematically performed *in silico* knockouts of individual TFs within each network. Using topic matrix loadings (Figure 1A), we constructed functional gradients for topics of interest, and calculated dot products reflecting the magnitude of alteration of these gradients as a consequence of each TF perturbation, resulting in topic-specific perturbation scores (Figure 2B). When homeostatic and inflammatory perturbation scores were projected as vectors in two-dimensional space, vectors corresponding to recruited AMs had a greater average magnitude than those from tissue resident AMs (Figure 2C). This finding is consistent with a relative lack of stabilizing regulatory interactions for recruited AMs, such that perturbation of a single TF produces a larger functional shift. These results support a model in which dense TF interconnectivity stabilizes GRN architecture and constrains inflammatory responses in tissue resident AMs, consistent with their primary role in maintaining lung homeostasis.

**Figure 2.**
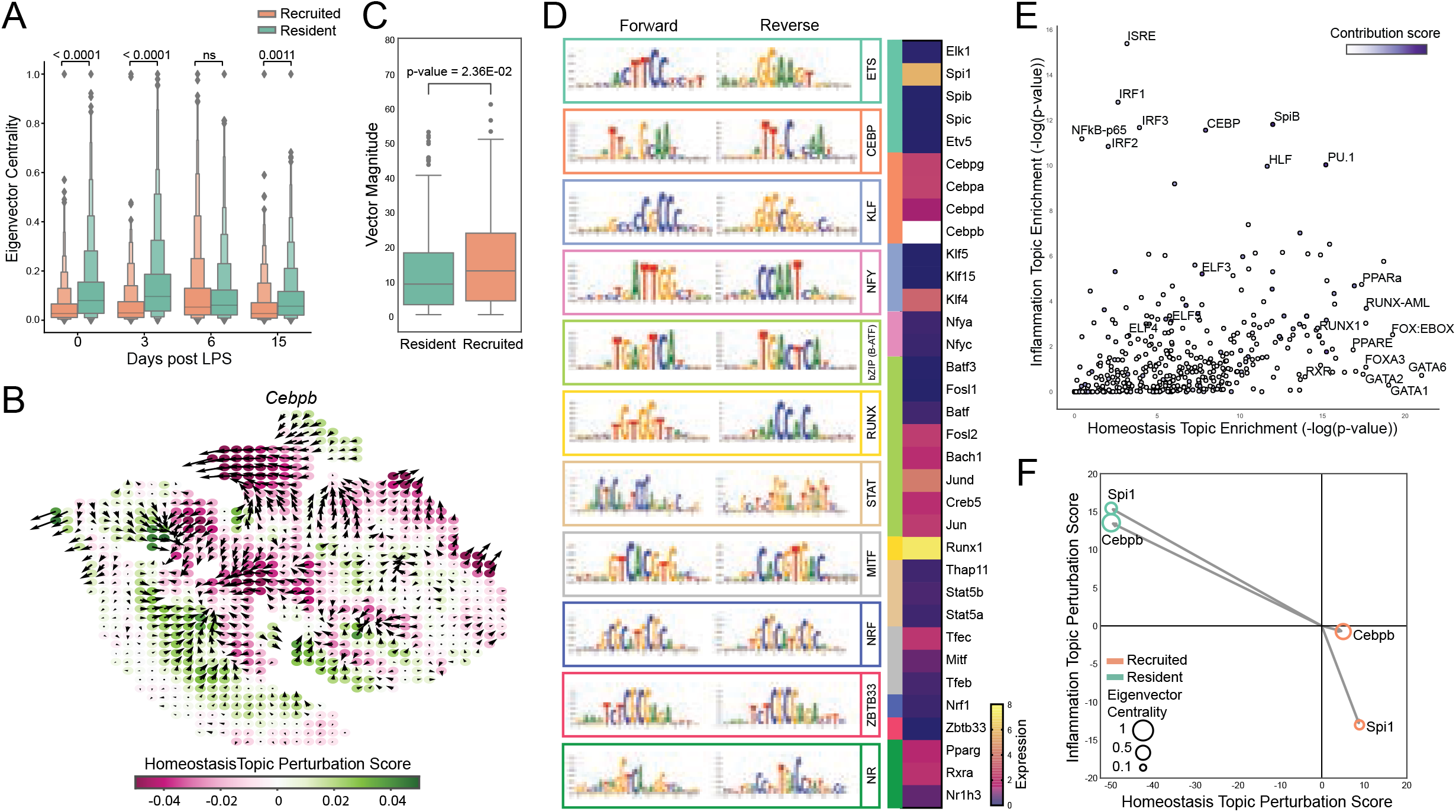
Transcription factors with high contribution to chromatin accessibility strongly influence the magnitude of functional shift after in silico perturbation. a) Boxenplots comparing average eigenvector centralities per AM type across the CellOracle gene regulatory networks for each timepoint (p-values are from 2-way ANOVA with multiple comparisons) b) Example perturbation simulation trajectories after *in silico* knockout of *Cebpb* and the perturbation scores with respect to the homeostasis topic trajectory. c) Box plot showing the average homeostatic by inflammatory perturbation score vector magnitudes by AM type for all TFs. d) De novo motif discovery from high contribution score regions from ChromBPNet as determined by TF-MoDISco. The forward and reverse motifs are given, as well as the transcription factor family matched by similarity. The basal normalized transcriptomic expression is given for transcription factors in these families (*Cebpb* expression is 23.7 and is shown in white). e) Scatterplot of HOMER-discovered motifs found in the promoter regions (within 100kbp of TSS) of genes differentially expressed in either inflammation topic (y-axis) or homeostasis topic (x-axis). Motifs are colored by their average contribution score determined by ChromBPNet. f) Vectors of net perturbation scores for the homeostasis topic (x-axis) and inflammation topic (y-axis) trajectories for *Cebpb* and *Spi1* for both AM types. The size of the circle corresponds to that TF’s eigenvector centrality in the GRN.

To further define the regulatory features controlling inflammatory responses in tissue-resident AMs, we generated ATAC-seq profiles of tissue-resident AMs (CD45^+^CD64^+^CD11b^−/lo^SiglecF^+^) isolated from unstimulated mice or mice three days after intratracheal LPS administration, and applied ChromBPNet(21) to identify sequence determinants of chromatin accessibility. Leveraging TF-MoDISco to cluster high-contribution genomic regions for *de novo* motif discovery, we found that the most abundant motifs in the accessible chromatin belonged to the ETS and CEBP families (Figure 2D), with *Spi1* and *Cebpb* emerging as the most highly expressed TF family members, consistent with their established roles in tissue-resident AM homeostasis. To relate chromatin features to gene expression programs, we performed HOMER motif analysis on open chromatin regions from both steady state and day 3 LPS conditions and assessed motif enrichment within 100 kb of transcription start sites for homeostasis and inflammation topic genes (Figure 2E). A diverse set of motifs were enriched near homeostasis topic genes, reflecting the involvement of diverse TFs in maintaining tissue-specific homeostatic functions of tissue-resident AMs. In contrast, motifs selectively enriched near inflammation topic genes included IRFs, ISRE, and NF-κB motifs, consistent with their well-characterized roles in macrophage inflammatory responses (22, 23). Notably, high-contribution motifs—including CEBP, PU.1, SpiB, and HLF (CEBP-related)—were enriched upstream of both homeostasis and inflammation topic genes. These findings suggest that high-contribution transcription factors CEBP/β and PU.1 may maintain the regulatory landscape that coordinates functional plasticity of tissue-resident AMs. To explore this possibility, we returned to our GRN analysis and examined the predicted impact of perturbing *Cebpb* (CEBP/β) and *Spi1* (PU.1) on homeostatic and inflammation topic expression. Strikingly, in silico knockout of these high contribution TFs in tissue-resident AMs produced large vector magnitudes, reflecting pronounced shifts away from homeostatic programs and toward inflammatory states (Figure 2F). These results suggest that CEBP/β and PU.1 promote homeostatic programming while constraining inflammatory programs in tissue-resident AMs. Notably, perturbations of these TFs yielded larger vector magnitudes in tissue-resident AMs than in recruited AMs (Figure 2F), opposite to our observation that on average, global TF perturbations had reduced impact in tissue-resident AMs (Figure 2C). Together, these results suggest that CEBP/β and PU.1 stabilize GRN architecture in tissue-resident AMs and constrain inflammatory gene expression through coordinated regulation of downstream targets.

We have systematically investigated TF regulatory interactions that govern AM functional plasticity in response to changes in the tissue environment. CEBP/β and PU.1 are well established as key regulators of tissue-specific homeostatic AM functions. The high contribution and network centrality of CEBP/β and PU.1 identified here suggest that these TFs are central to maintaining tissue-resident AM identity in response to environmental stimuli. Given evidence that these TFs can act as pioneer factors and cooperatively bind at regulatory regions with other TFs(24–27), we speculate that their stabilizing influence is supported by TF cooperation at the chromatin level that underlies tissue-specific AM responses to inflammatory stimuli. This model is consistent with the markedly distinct chromatin landscape of recently recruited AMs compared to tissue-resident AMs, which may permit recruited AMs to sustain a pro-inflammatory transcriptional profile upon exposure to the same lung environment (15). More broadly, this integrative framework combining tissue-specific GRNs with chromatin accessibility analysis can be applied to various tissue contexts to understand transcriptional mechanisms underlying healthy and dysregulated macrophage functional states.

## Materials and Methods

### Data Processing

Single-cell RNA-seq data was acquired from GSE280003, which was then filtered to obtain only the gene expression values from the sorted alveolar macrophages. K-means clustering of this data allowed for the identification of two main clusters out of five that best represented tissue-resident and monocyte-derived macrophages, and was used to further filter the data to analyze only these cells. For pseudobulk data, the average expression of each gene across origin and/or timepoint was taken at each time point. For ATAC-seq processing, adapters were trimmed from the fastq files before aligning to the mm10 genome. After sorting and removing duplicate reads, peaks were called using MACS2 at a q-value of 0.01. The intersection of the peaks was taken between biological replicates for further analysis.

### Topic Modeling

Topic modeling was performed via the R package fastTopics. In brief, a non-negative matrix factorization was applied to the single cell expression counts and was reparametrized to reflect the proportion of each topic to the counts and the probability of a given cell belonging to each topic. Model fitting was performed over 150 iterations with an additional 150 iterations to fine-tune. Models were evaluated for best convergence to a state of maximum likelihood and minimizing residuals. We selected a model with k=7 total topics for best model performance. Using grade-of-membership differential expression analysis(17), we took the top 1000 genes with the greatest log fold change for KEGG pathway enrichment analysis and used this to identify 3 topics that related to macrophage functional states to investigate further. The remaining topics either had no clear functional annotation or were determined to be driven by contaminating cells and were not analyzed further.

### ATAC-seq and Contribution Scores

Samples and data for ATAC-seq were prepared as previously described (28). Briefly, AMs from naïve and LPS-treated mice were sorted by FACS, from which DNA was isolated and amplified with indexed primers. The bar-coded amplicons were sequenced in a NovaSeq SP (Illumina) under a 2×150 pair-ended format and fastq files were generated per the standard Illumina pipeline. Adapter sequences were trimmed from the reads and aligned to the mm10 reference genome with Bowtie2 and filtered for only high-quality uniquely mapped reads. Peaks were called with MACS2 with default parameters and peaks were filtered based on FDR <0.01. Peaks were merged across into consensus peak regions with BEDTools and filtered out mm10 blacklisted regions from ENCODE. ATAC-seq data from GSE100738 was processed as described and used to train alveolar macrophage ChromBPNet models, including a custom Tn5-bias model to regress out Tn5 cut bias from ATAC-seq sample processing. Five separate models were trained on five separate folds, which were each comprised of different combinations of training, validation, and testing chromosome groups. These models were used to calculate the contribution scores of open chromatin regions from naïve basal AMs isolated from control mice and from AMs isolated from mice that were day 3 after LPS exposure (GSE231217, GSE319365), and the contribution score bigwigs were merged for each sample across all folds.

### Motif Analysis

For de novo motif analysis, we utilized TFModisco from the python package modisco-lite on the basal and 3-day LPS counts and profile contribution scores. Motif patterns were manually annotated to TF families based on significant matches. For motif hit calls of known motifs, we utilized HOMER motif finding software on the union set of peaks from the naïve and LPS-treated AM ATAC-seq data. With the resulting bed file, we used the R package ChIPseeker to annotate these motifs to putative target genes within 3kb from their transcription start site. For the average contribution scores of these motifs, we use bigwigaverageOverBed to calculate the average contribution score using the counts contribution bigwig file for the naïve condition over each motif region, and then averaged those contribution score averages.

### Gene Regulatory Network Construction

To generate gene regulatory networks using CellOracle, we generated a base GRN using the Tn5 bias-corrected open chromatin regions from the ChromBPNet model using CellOracle’s pipeline, which utilizes the gimmemotifs python package for motif scanning. Motifs were scanned with a background of 200bp and were then filtered for scores of 10 or greater. The scRNA-seq data was filtered for at least one expressed gene and then further filtered for the top 2500 most variable genes. A list of expressed TFs analyzed by proteomics in AMs(29) was appended to this list of genes for a total of 2850 genes for network construction. KNN imputation was performed with the first 47 principle components and k = 598 (2.5% of the number of cells) as part of the network construction pipeline.

### Systematic in silico Perturbation

Within the CellOracle framework, we created gradients using the topic model embeddings to establish topic trajectories for topics k4, k5, and k7 for both tissue-resident and monocyte-derived populations. We then systematically perturbed *in silico* each transcription factor by way of simulating a knockout by reducing the expression to 0. For each in silico knockout, the dot products were calculated from the resulting perturbation trajectory and each topic trajectory and considered as perturbation scores representing the influence of that transcription factor in maintaining the predefined functional state (topic).

## Acknowledgments

We would like to thank members of the Gottschalk lab and Center for Systems Immunology for constructive discussion. Specifically, we thank Peter Gerges, Nicolas Pease, and Swapnil Keshari for advice and sharing methods related to the ChromPBnet workflow, and Hanxi Xiao and Tracy Tabib for assistance with CellOracle. We also thank the Center for Research Computing for providing computing resources (this work used the HTC and H2P clusters supported by NIH award S10OD028483 and NSF award OAC-2117681, respectively).

